# Genetic background affects the strength of crossover interference in house mice

**DOI:** 10.1101/2024.05.28.596233

**Authors:** Andrew P Morgan, Bret A Payseur

## Abstract

Meiotic recombination is required for faithful chromosome segregation in most sexually reproducing organisms and shapes the distribution of genetic variation in populations. Both the overall rate and the spatial distribution of crossovers vary within and between species. Adjacent crossovers on the same chromosome tend to be spaced more evenly than expected at random, a phenomenon known as crossover interference. Although interference has been observed in many taxa, the factors that influence the strength of interference are not well understood. We used house mice (*Mus musculus*), a well-established model system for understanding recombination, to study the effects of genetics and age on recombination rate and interference in the male germline. We analyzed crossover positions in 503 progeny from reciprocal F1 hybrids between inbred strains representing the three major subspecies of house mice. Consistent with previous studies, autosomal alleles from *M. m. musculus* tend to increase recombination rate, while inheriting a *M. m. musculus* X chromosome decreases recombination rate. Old males transmit an average of 0.6 more crossovers per meiosis (5.0%) than young males, though the effect varies across genetic backgrounds. We show that the strength of crossover interference depends on genotype, providing a rare demonstration that interference evolves over short timescales. Differences between reciprocal F1s suggest that X-linked factors modulate the strength of interference. Our findings motivate additional comparisons of interference among recently diverged species and further examination of the role of paternal age in determining the number and positioning of crossovers.

## Introduction

Meiotic recombination is a potent cellular and evolutionary force. In most species that reproduce sexually, recombination is required for the segregation of chromosomes during meiosis, with too few or too many crossovers leading to aneuploid offspring that are usually either non-viable or phenotypically abnormal (reviewed in Hassold and Hunt (2001)). Recombination creates new allelic combinations and thereby shapes the trajectories of beneficial and deleterious mutations in populations (Fisher 1930; Hill and Robertson 1966; Felsenstein 1974; Barton 1995).

Although its functional and evolutionary roles impose constraints on recombination, the number and positioning of crossovers nonetheless vary among individuals, both within and between species (Smukowski and Noor 2011; Stapley *et al*. 2017; Brazier and Glémin 2022). Evidence for a heritable component to this variation comes from responses to artificial selection (Chinnici 1971; Kidwell and Kidwell 1976), resemblance among relatives (Kong *et al*. 2004; Coop *et al*. 2008; Brekke *et al*. 2022), and the identification of genetic variants associated with recombination rate differences in populations (Kong *et al*. 2008; Johnston *et al*. 2016; Hunter *et al*. 2016). Likewise, the fine-scale spatial distribution of recombination has been shown to vary within and between populations in several taxa (reviewed in Paigen and Petkov (2010)). In mammals, one determinant of this variation is the interaction between by nucleotide sequence at so-called “hotspots” and the protein PRDM9, which applies chromatin marks that designate the location of double-strand breaks that may mature into crossovers (Parvanov *et al*. 2010).

Despite ample evidence for genetic variation in the overall rate and fine-scale distribution of recombination, the causes of the relative positions of crossovers on the same chromosome at macroscopic scale remain poorly understood. Crossovers are spaced more evenly than expected if they were positioned at random with respect to each other, an observation made during the construction of the first linkage maps (Sturtevant 1913, 1915). This pattern implies that the designation of a crossover in one chromosomal interval inhibits the formation of another crossover nearby, a phenomenon known as crossover interference. Crossover interference is observed across a wide variety of species (reviewed in Berchowitz and Copenhaver (2010); Otto and Payseur (2019)), with *Schizosaccharomyces pombe* (Munz 1994), *Aspergillus nidulans* (Strickland 1958), and *Aspergillus fumigatus* (Auxier *et al*. 2023) providing documented exceptions.

The magnitude of crossover interference is rarely compared across taxa, but distantly related species appear to differ (Otto and Payseur 2019). For example, interference in dogs is estimated to be 2 *—* 3 times as strong as interference in humans (Campbell *et al*. 2016). The main evidence for variation in interference within species comes from contrasts between the sexes (Otto and Payseur 2019). Males show stronger interference in humans (Campbell *et al*. 2015), dogs (Campbell *et al*. 2016), mice (Peterson and Payseur 2021), and gray geese (Torgasheva and Borodin 2017), whereas females show stronger interference in cattle (Wang *et al*. 2016) and *Arabidopsis thaliana* (Basu-Roy *et al*. 2013). In *A. thaliana*, a lower proportion of crossovers escapes interference in females than in males (Basu-Roy *et al*. 2013).

The extent to which the magnitude of interference and the percentage of interfering crossovers vary among individuals of the same sex is virtually unknown. One study in cattle found an association between the strength of interference in males and variation in *Nek9*, a gene involved in meiotic spindle formation, implying the existence of heritable differences (Wang *et al*. 2016). A quantitative trait locus modulates a difference in interference between two wild-derived strains of house mice (Wang *et al*. 2019). The absence of information about variation in interference among individuals is striking given that this phenomenon is a key determinant of crossover positioning and exerts substantial influence on the distribution of relatedness in populations (Caballero *et al*. 2019; Veller *et al*. 2020).

House mice from the *Mus musculus* species complex provide a useful system for examining variation in crossover interference. Interference in mice is strong, with distances between double crossovers that amount to appreciable percentages of chromosomal length (Broman *et al*. 2002). For example, the minimal physical separation between double crossovers on chromosome 1 detected in 3 026 male mouse meioses was 57 Mb (Petkov *et al*. 2007; Paigen *et al*. 2008). Variation in crossover interference has been proposed to explain sex differences in recombination rate in mice (Petkov *et al*. 2007). The molecular and cellular pathways that culminate in crossovers are well understood in mice (reviewed in Lam and Keeney (2014) and Gray and Cohen (2016)). Furthermore, wild-derived inbred strains capture natural variation in recombination rate (Peterson and Payseur 2021) and enable reproducible comparisons of interference. Here, we examine the transmission of crossovers in F1 hybrids generated from crosses between wild-derived strains of mice belonging to the three principal subspecies of house mice: *M. m. domesticus*, *M. m. musculus* and *M. m. castaneus*. We uncover heterogeneity in the strength of crossover interference, demonstrating that this important determinant of the recombination landscape evolves on short timescales.

## Materials and methods

### Mouse breeding

Mice were generated for this study as part of a larger research program studying the effects of aging on reproductive phenotypes in hybrid males. The breeding scheme used in this work has been described previously (Morgan *et al*. 2019) and is shown in **Figure 1**. Reciprocal F1 hybrid males were bred in all possible pairwise combinations between CAST/EiJ (*M. m. castaneus*), PWK/PhJ (*M. m. musculus*) and WSB/EiJ (*M. m. domesticus*). Founder individuals were purchased from the Jackson Laboratory (Bar Harbor, ME) between 2008 and 2011. Shortly after reaching sexual maturity, F1 males were mated to multiple FVB/NJ females and liveborn N2 progeny were collected and sacrificed at birth by cervical dislocation. At least 2 litters were collected from each male by age 4 months (“young age”). F1 males were allowed to continue mating freely, and females were replaced if they stopped producing offspring. Once F1 males were approaching reproductive senescence (as evidenced by increasing interval between litters or declining litter size), at least 2 additional litters were collected at an age of at least 15 months (“old age”). Breeding was completed in June 2011. The number of F1 males used, progeny collected, and proportion used for genotyping are shown in **Table 1**. The distribution of litter sizes for each F1 genotype is shown in **Figure S1**. The list of N2 progeny used in this study is provided in **Table S1**.

**Figure 1:**
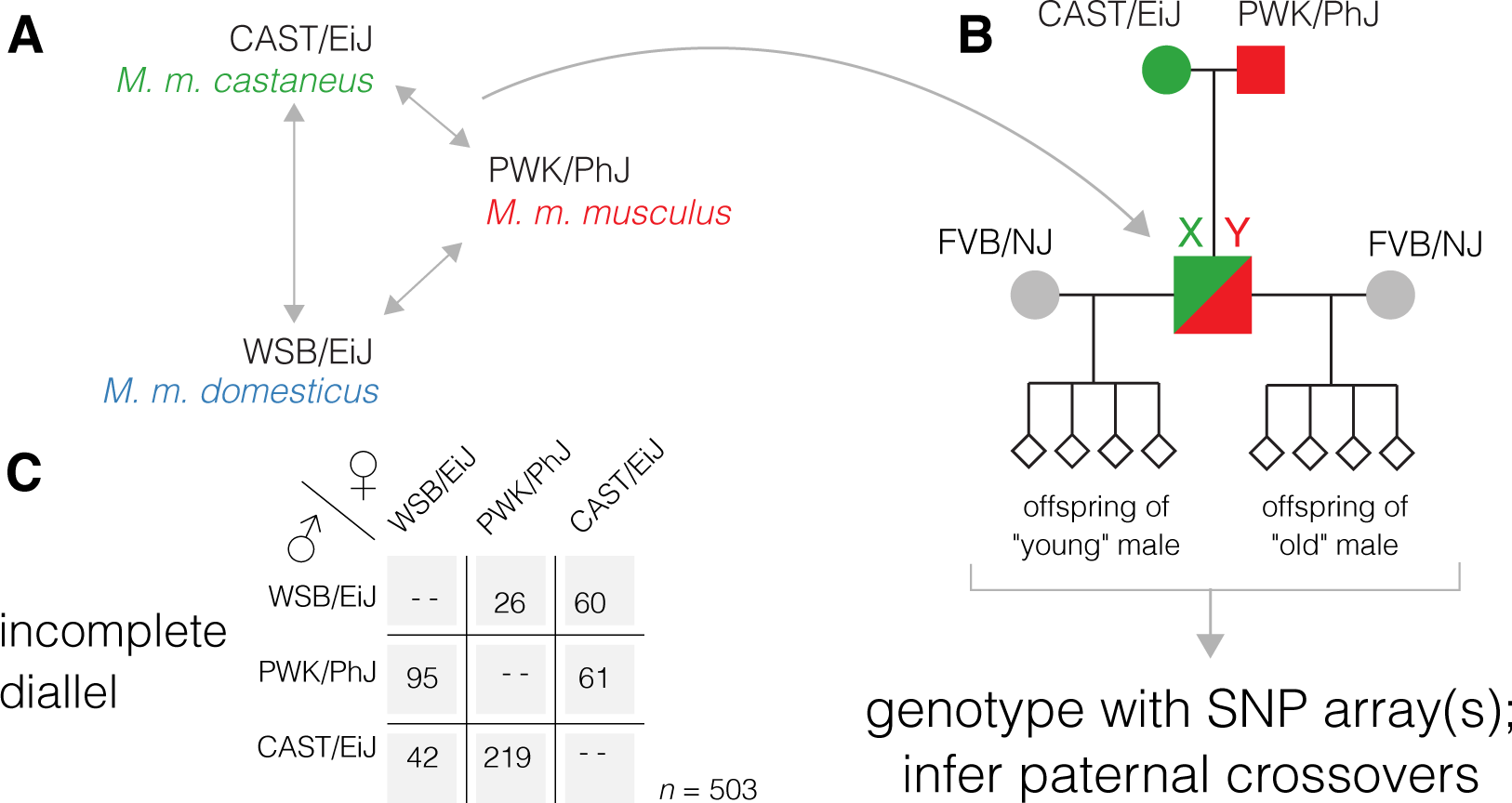
Incomplete diallel design to study effects of age and genetics on recombination rate. (**A**) All possible F1 genotypes between three parental strains representing the cardinal subspecies of *Mus musculus*. (**B**) F1 males mated at young age (< 4 months) and old age (>15 months) to highly fertile FVB/NJ females. (**C**) Number of progeny collected for each paternal genotype.

**Table 1:** Number of breeding males, total progeny, and progeny used by genotype.

| genotype | # breeding males | # progeny | proportion used |
| --- | --- | --- | --- |
| CAST/EiJ × PWK/PhJ | 2 | 98 | 0.51 |
| PWK/PhJ × CAST/EiJ | 5 | 259 | 0.67 |
| CAST/EiJ × WSB/EiJ | 4 | 234 | 0.15 |
| WSB/EiJ × CAST/EiJ | 4 | 258 | 0.28 |
| PWK/PhJ × WSB/EiJ | 4 | 80 | 0.48 |
| WSB/EiJ × PWK/PhJ | 6 | 329 | 0.29 |

Although the number of progeny actually used from each genotype does vary widely (range 26 *—* 219), this reflects decisions about resource allocation to balance competing experimental priorities unrelated to the work described in this manuscript, and is not solely attributable to differences in fertility. The study was approved by the Institutional Animal Care and Use Committee of the University of North Carolina at Chapel Hill, and all animal husbandry was conducted in the university’s Association for Assessment and Accreditation of Laboratory Animal Care-accredited facility (#A3410-01; Animal Welfare Assurance) in accordance with institutional and federal regulations.

### DNA extraction and genotyping

As previously described (Morgan *et al*. 2019), genomic DNA was extracted from whole tails or heads of N2 mice and from tail clips of adult representatives of parental inbred strains, using DNeasy Blood & Tissue Kits (catalog no. 69506; QIAGEN, Valencia, CA). Approximately 1.5 *µ*g of DNA per sample was shipped to Neogen (Lincoln, NE) for genotyping with one of two Mouse Universal Genotyping Arrays, dubbed MegaMUGA (approximately 77K probes) (Morgan *et al*. 2016) or MiniMUGA (approximately 11K probes) (Sigmon *et al*. 2020). Both are designed on the Illumina Infinium HD platform (Steemers *et al*. 2006) and SNP content is optimized for genetic mapping in populations descended from common laboratory and wild-derived mouse strains. (The MegaMUGA product was discontinued partway through the study, so remaining genotyping used MiniMUGA.) The vendor provided raw genotype calls at all markers on each array. The number of N2 progeny genotyped on each array, from each genotype and paternal age, are shown in **Table 2**.

**Table 2:**
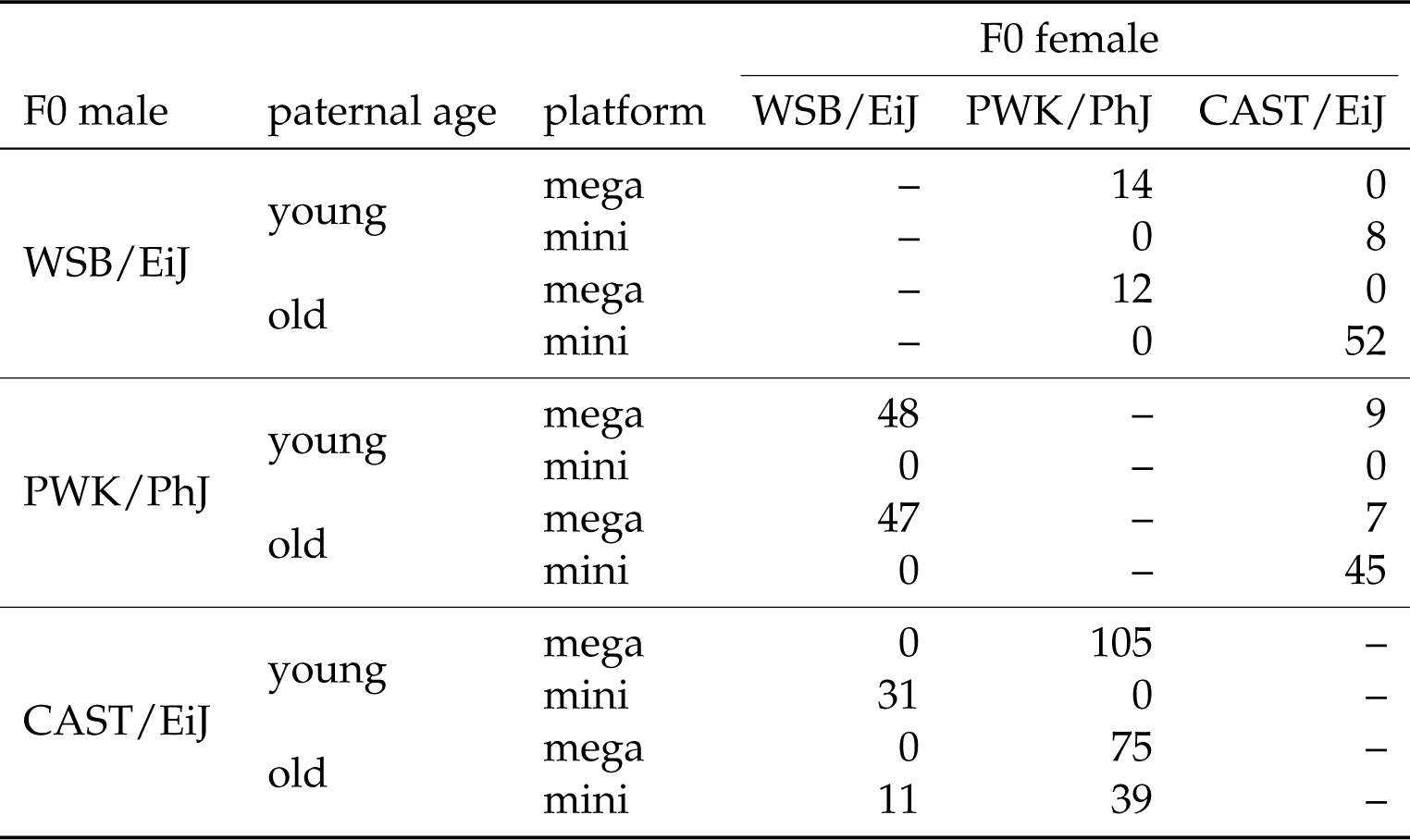
Counts of N2 progeny by genotype, paternal age, and genotyping platform (“mega” = MegaMUGA, “mini” = MiniMUGA).

| F0 male | paternal age | platform | F0 female |  |  |
| --- | --- | --- | --- | --- | --- |
|  |  |  | WSB/EiJ | PWK/PhJ | CAST/EiJ |
| WSB/EiJ | young | mega | – | 14 | 0 |
|  |  | mini | – | 0 | 8 |
|  | old | mega | – | 12 | 0 |
|  |  | mini | – | 0 | 52 |
| PWK/PhJ | young | mega | 48 | – | 9 |
|  |  | mini | 0 | – | 0 |
|  | old | mega | 47 | – | 7 |
|  |  | mini | 0 | – | 45 |
| CAST/EiJ | young | mega | 0 | 105 | – |
|  |  | mini | 31 | 0 | – |
|  | old | mega | 0 | 75 | – |
|  |  | mini | 11 | 39 | – |

### Filtering and harmonizing genotypes

Genotype data were processed in `R` v4.1.1 (R Foundation for Statistical Computing, http://www.r-project.org/) using the `argyle` package v0.2.2 (Morgan 2016). Physical positions (in the GRCm39 assembly) and male-specific genetic positions on the standard mouse genetic map (Cox *et al*. 2009) for array markers were assigned based on the annotations produced by Karl Broman and colleagues (https://github.com/kbroman/MUGAarrays). Initial filtering and quality control steps were performed separately for each combination of array and pater-nal genotype, since different subsets of markers will be informative depending on paternal genotype. Autosomal markers which met the following criteria were retained: (1) *<* 10% missing genotypes; (2) fixed for opposite alleles in the founder strains; and (3) not subject to substantial segregation distortion (non-reference allele frequency *>* 20% and *<* 80%). For all paternal genotypes besides PWK/PhJ *⇥* WSB/EiJ and WSB/EiJ *⇥* PWK/PhJ, genotyping was split between the two array platforms. To mitigate biases arising from differential marker density, for each marker on the more sparse array (MiniMUGA), the nearest marker on the more dense array (MegaMUGA) was retained, and other markers were discarded. (This approach retains markers that are shared between the two arrays.) We also sought to mitigate technical biases arising from differences between array platform in the proportion of each chromosome spanned by informative markers, as follows. For each paternal genotype, let (*p*_mini_*, d*_mini_) and (*p*_mega_*, d*_mega_) be the most proximal and distal markers on MiniMUGA and MegaMUGA, respectively; and let *p*_inner_ = *max p*_mini_*, p*_mega_ and *d*_inner_ = *min d*_mini_*, d*_mega_ . Any markers proximal to *p*_inner_ or distal to *d*_inner_ were discarded. The number of markers retained for each combination of paternal genotype and platform, and their physical span, are shown in **Table 3**.

### Haplotype and crossover inference

Filtered genotypes were converted to `R/qtl2` format, recoded with respect to the inbred paternal grandparents. Although our breeding scheme is not actually a backcross, expected genotype frequencies are 0.5 in N2 offspring so genotype probabilities were modelled as a a backcross using `R/qtl2`’s hidden Markov model (HMM) (Broman *et al*. 2019). The most likely sequence of haplotype states at each marker was inferred using the Viterbi algorithm (`qtl2::viterbi()`), allowing 1% genotyping error. Uninterrupted runs of the same marker-wise haplotype state thus correspond to transmitted haplotype segments, and crossovers are implicitly defined as boundaries between haplotype segments. Inferred haplotype segments are provided in **Table S2**, and the number of autosomal crossovers per individual in **Table S3**.

**Table 3:** Density and coverage of pruned marker maps by genotype and platform (“mega” = MegaMUGA, “mini” = MiniMUGA, *n* = number of N2 progeny genotyped).

| genotype | platform | markers | Mb covered | $n$ |
| --- | --- | --- | --- | --- |
| CASTxPWK | mega | 1940 | 2352 | 16 |
|  | mini | 2059 | 2354 | 45 |
| PWKxCAST | mega | 1940 | 2352 | 180 |
|  | mini | 2059 | 2354 | 39 |
| CASTxWSB | mega | – | – | 0 |
|  | mini | 2712 | 2359 | 60 |
| WSBxCAST | mega | – | – | 0 |
|  | mini | 2712 | 2359 | 42 |
| PWKxWSB | mega | 32010 | 2393 | 26 |
|  | mini | – | – | 0 |
| WSBxPWK | mega | 32005 | 2393 | 95 |
|  | mini | – | – | 0 |

### Statistical analyses of recombination rate

Genetic and non-genetic effects on the autosome-wide (“global”) recombination rate were modelled by treating our experiment as an incomplete diallel between WSB/EiJ, PWK/PhJ and CAST/EiJ. The phenotype of interest is the number of autosomal crossovers per meiosis, and each N2 offspring represents an independent observation. Additive, parent-of-origin, strain-pair-specific, and age effects were estimated using the `BayesDiallel` package v0.982 (Lenarcic *et al*. 2012). The hierarchical Bayesian model implemented in this package can also include terms for inbreeding and dominance effects, but without phenotypes measured in inbred parental strains, these cannot be estimated. The Gibbs sampler was run for 10 000 iterations (the first 1 000 of which were discarded as burn-in), in 3 replicate chains.

### Statistical analyses of crossover interference

Crossover interference was analyzed under two well-studied models: the gamma model, in which the strength of interference is expressed by a single unitless parameter, *ν* (Broman and Weber 2000); and the gamma-escape model of Housworth and Stahl (2003), in which a proportion *p* crossovers are not subject to interference, and the remaining 1 *— p* are subject to interference under the gamma model with parameter *ν*. We sought genotype-specific estimates of the strength of crossover interference, and allowed this effect to vary with paternal age. First we fit the gamma and gamma-escape models to each subgroup (defined by paternal genotype, or the combination of paternal genotype and paternal age cohort) separately, using the maximum-likelihood approach implemented in the `xoi` package v0.72 (Broman 2023). Uncertainty in parameter estimates was obtained by performing 1000 bootstrap replicates, in which individuals were sampled randomly with replacement from within each group. This approach implicitly assumes no relationship between group-specific parameter values. We reasoned that the strength of interference is likely biologically constrained and that treating group-specific parameter values as being drawn from a common distribution would provide regularization that might lead to more stable and precise estimates. The following hierarchical Bayesian model was therefore implemented. Let *ν_i_* be the group-wise interference, and define it as follows:

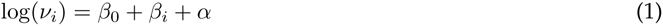

and define the following terms:

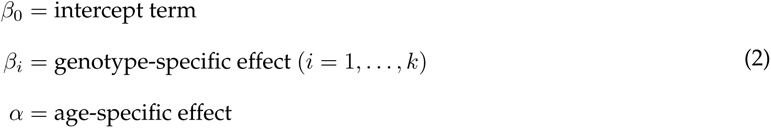

The prior distributions for these parameters were *β*_0_ *∼ N* (*µ, σ*^2^), and the remaining *β_i_ ∼ N* (0*, σ*^2^). The prior mean (*µ* = 1) and variance (*σ*^2^ = 0.5) were chosen arbitrarily, but we experimented with a range of hyperparameter values and found they had little impact on results. Similarly:

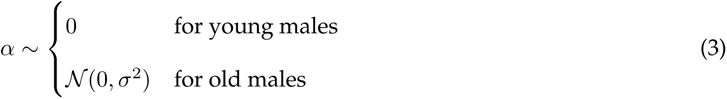

We implemented a simple Metropolis-coupled Markov chain Monte Carlo (MCMC) sampler to do inference of these parameters. The proposal distribution was *N* (0*, τ* ^2^), and we found that *τ* ^2^ = 0.005 allowed adequate mixing. The likelihood of a parameter vector *θ* = (*β*_0_*,. .., β_k_, λ*) given the complete set of haplotype segments was calculated with the `C` routine `GammaS()`, which is exposed in the `xoi` package. The sampler was run for 25 000 iterations (the first 5 000 of which were discarded as burn-in), in 4 replicate chains. Time required to compute the likelihood grows approximately linearly in the number of haplotype segments, so runtime of each MCMC chain was about one day. Convergence and mixing were assessed by manual inspection of trace plots and by Gelman-Rubin diagnostics as implemented in the `coda` package (Plummer *et al*. 2006). Out-of-sample predictive performance was assessed using smoothed importance sampling as implemented in the `loo` package, and influential outliers were defined as those with Pareto *k* scores *≥* 0.5 (Vehtari *et al*. 2017). The UNC Research Computing cluster was used for most computation.

## Results

### Recombination rate varies by genotype and age

The study design is shown in **Figure 1**. A total of 503 offspring of 17 males representing 6 reciprocal F1 crosses were collected and genotyped with SNP arrays. After iterative filtering and pruning, marker maps cover at least 97.8% of the physical span of the autosomes in each paternal genotype. The overall genotyping rate was 99.8%; no individual had more than 1.2% genotypes missing. We identified a total 15 744 haplotype segments that define 6 288 autosomal crossovers. On average 12.3 crossovers (range 5 *—* 22) were transmitted in each male meiosis. Consistent with many prior studies, crossovers were concentrated in the distal ends of chromosomes (**Figure S2**).

The distributions of crossover count by genotype and paternal age are shown in **Figure 2**. The effects of interest are the influence of genotype and paternal age on recombination rate. As a preliminary step, we analyzed crossover counts under a generalized linear model (with Poisson response and log link) with terms for genotype, age, and genotype-by-age interaction. There is significant heterogeneity in recombination rate across genotypes (*p* = 2.5 *×* 10*^—^*^9^, likelihood ratio test). Qualitatively, paternal age appears to have a modest effect that differs by genotype. In a model allowing an age-by-genotype interaction, neither the main effect of age nor its interaction term reach statistical significance (**Table 4**). However, treating observations as conditionally independent, given genotype, ignores correlation structure that may be present when genotypes share parents. We therefore re-analyzed our experiment as an incomplete diallel between WSB/EiJ, PWK/PhJ and CAST/EiJ. Additive, parent-of-origin and cross-specific genetic effects were included in the model. Results are shown in **Figure 3**. Alleles from the PWK/PhJ background tend to increase the recombination rate, but having a PWK/PhJ mother decreases recombination rate by 0.51 (95% highest posterior density interval [HPDI] 0.28 *—* 0.74) crossovers per meiosis. In this analysis, older males transmit 0.61 (95% HPDI 0.08 *—* 1.13) more crossovers than younger males. These effects are modest: the contributions of additive, parent-of-origin and age effects are 15.1%, 8.2% and 0.8% of trait variance, respectively.

**Figure 2:**
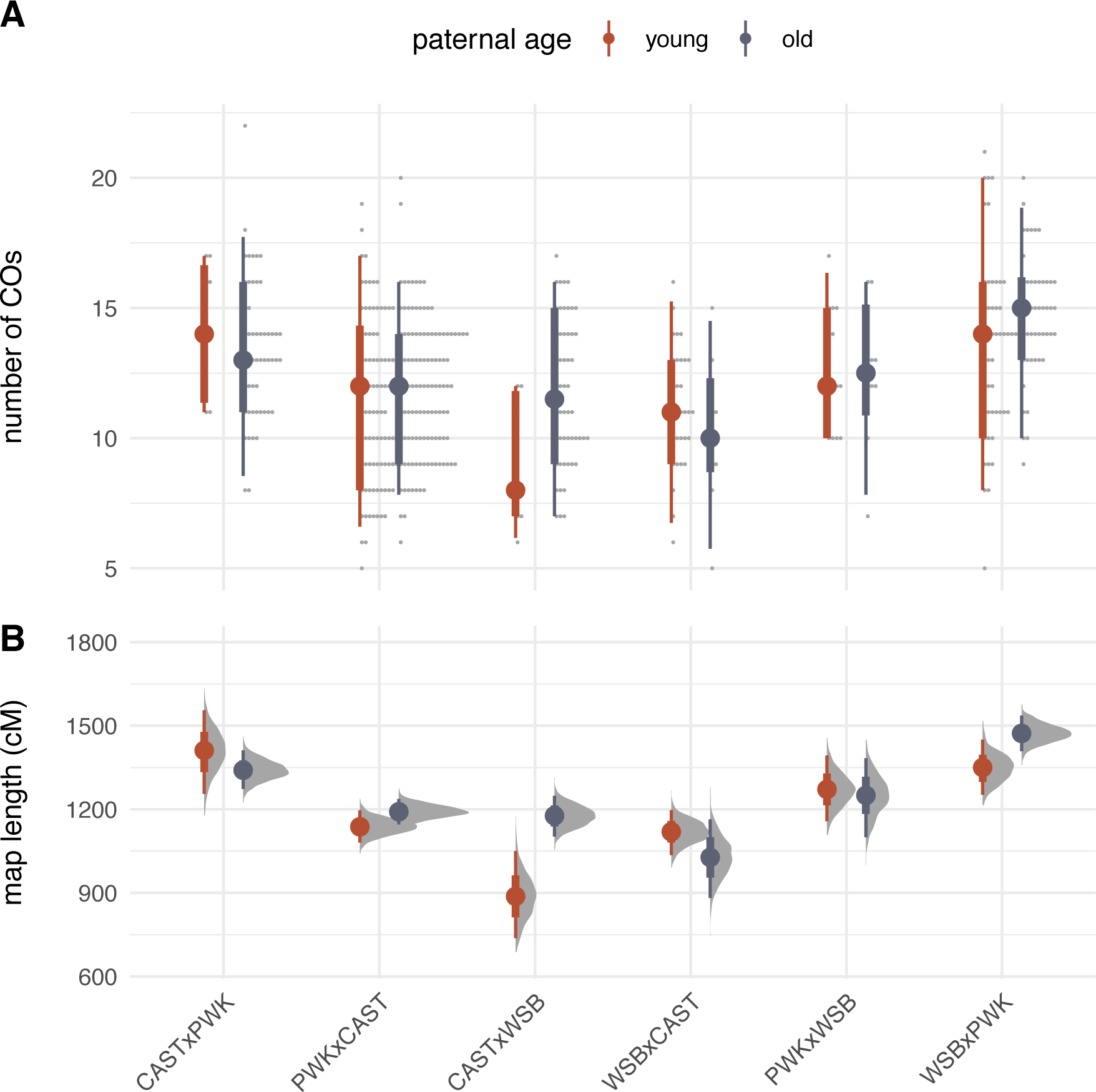
Recombination rate by genotype and age. (**A**) Number of transmitted crossovers per meiosis. Each point is an individual observation; solid dots show group means and bars show 2.5% – 97.5% (thin) and 25% – 75% (thick) quantile intervals. (**B**) Bootstrap estimates of the length of the autosomal recombination map in each genotype. Solid dots show group means; error bars defined as in panel **A**.

**Figure 3:**
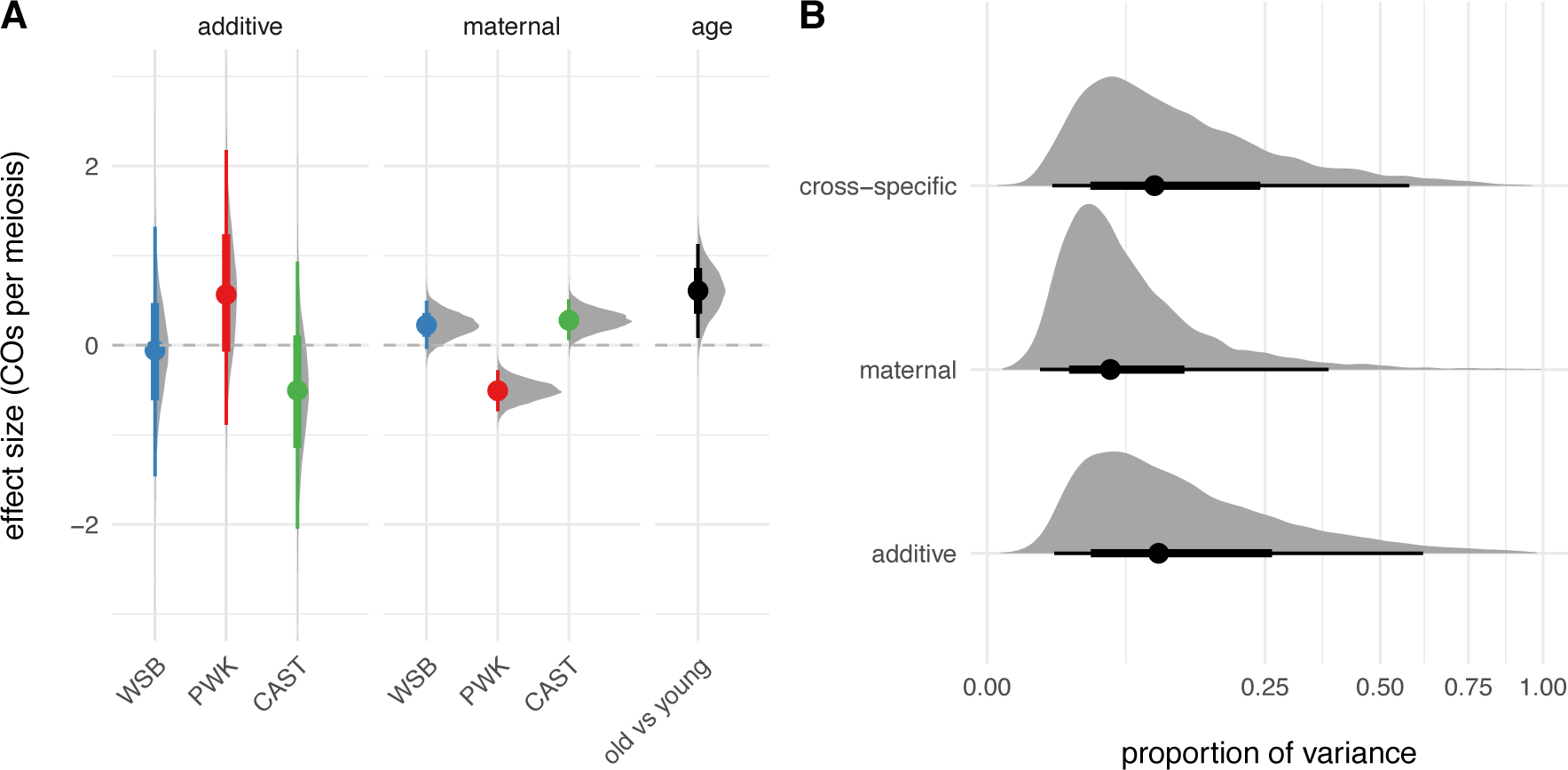
Diallel analysis of recombination rate. (**A**) Posterior distribution of estimated additive and maternal genotype effects (see main text) and age effect. Solid dots show group means and bars show 2.5% – 97.5% (thin) and 25% – 75% (thick) quantile intervals. (**B**) Posterior distribution of trait variance attributed to each class of genetic effects. Note square-root scale on horizontal axis. Quantile intervals indicated as in panel **A**.

**Table 4:** Analysis of deviance for model of crossover counts. The *p*-values shown are from a likelihood-ratio test. df = degrees of freedom.

| | df | deviance | residual df | residual deviance | $p$ |
| --- | --- | --- | --- | --- | --- |
| genotype | 5 | 48.72 | 497 | 320.25 | $2.5 \times 10^{-9}$ |
| age | 1 | 3.23 | 496 | 317.02 | 0.07 |
| genotype × age | 5 | 7.24 | 491 | 309.78 | 0.20 |
| residuals | — | — | 502 | 368.97 | — |

As expected, the number of chromosomes with multiple crossovers increases linearly with the total number of crossovers (**Figure S3**) . The distribution of inter-crossover distances on chromosomes with multiple crossovers is summarized in **Table 5**, and plotted against the expected distribution in the absence of interference in **Figure S4**. Qualitatively, the distribution of inter-crossover distances strongly suggests the presence of interference, and appears to differ between genotypes. We explore this possibility more rigorously in the next section. We find that inter-crossover distances appear to be shorter – implying weaker interference – on shorter chromosomes (**Figure S5**). This result contrasts with the pattern seen in female meiosis in previous work on *Mus musculus×Mus spretus* hybrids (Broman *et al*. 2002), but we did not attempt to explicitly estimate strength of interference on individual chromosomes.

**Table 5:** Distribution of inter-crossover distances (in cM) by genotype.

| genotype | mean | median | 2.5%ile | 25%ile | 75%ile | 97.5%ile |
| --- | --- | --- | --- | --- | --- | --- |
| CAST/EiJ × PWK/PhJ | 49.3 | 48.9 | 25.3 | 38.5 | 58.5 | 76.6 |
| PWK/PhJ × CAST/EiJ | 52.1 | 51.3 | 24.8 | 44.0 | 61.5 | 80.2 |
| CAST/EiJ × WSB/EiJ | 52.5 | 53.3 | 25.3 | 42.3 | 63.5 | 79.7 |
| WSB/EiJ × CAST/EiJ | 59.0 | 58.9 | 33.2 | 49.9 | 68.7 | 79.7 |
| PWK/PhJ × WSB/EiJ | 49.3 | 48.1 | 29.6 | 41.1 | 58.0 | 75.0 |
| WSB/EiJ × PWK/PhJ | 51.2 | 50.6 | 22.5 | 42.0 | 61.4 | 79.3 |

### Strength of crossover interference and interference escape vary by genotype

Crossover interference across all autosomes was quantified under two models: the gamma model (McPeek and Speed 1995; Broman and Weber 2000) and the Housworth-Stahl, or gamma-escape, model (Housworth and Stahl 2003). The gamma model describes the strength of interference via a single unitless parameter, *ν*; the distribution of intercrossover distances in this model has mean 50 cM and standard deviation 50*/√ν* cM. Although the gamma model provides a succinct statistical description of intercrossover distances, it is known that some proportion of crossovers are not subject to interference (reviewed by Gray and Cohen (2016).) The gamma-escape model allows a proportion *p* of non-interfering crossovers, with the remaining 1 *— p* subject to the gamma model of interference with strength *ν*. When *p* = 0, the gamma-escape model reduces to the gamma model, so the models are nested, allowing for model comparison via likelihood ratio test.

We used maximum likelihood to estimate (*ν_i_, p_i_*) separately for each genotype (for each of *i* = 0*,. ..k* genotypes) (full model), and then estimated a single pair of (*ν*_0_*, p*_0_) for the entire dataset (null model). Confidence bounds were obtained via nonparametric bootstrap. Results are shown in **Figure 4**. To evaluate the strength of evidence for heterogeneity of crossover interference across genotypes, we performed a likelihood ratio test between the full and null gamma models (restricting the *p_i_* = 0). Results are shown in **Table 6** and **Table 7**. The full gamma model is superior to the null (*p* = 0.0082), at the expense of increased model complexity. Allowing for interference escape that also varies across genotypes (comparing full gamma to full gamma-escape models) provides further improvement in fit (*p* = 2.9 *×* 10*^—^*^13^). We conclude that strength of crossover interference varies by genotype, and modelling the effect of interference escape improves fit to the observed distribution of haplotype segment lengths. Rate of interference escape appears to be positively correlated with strength of interference (**Figure 4C**).

**Figure 4:**
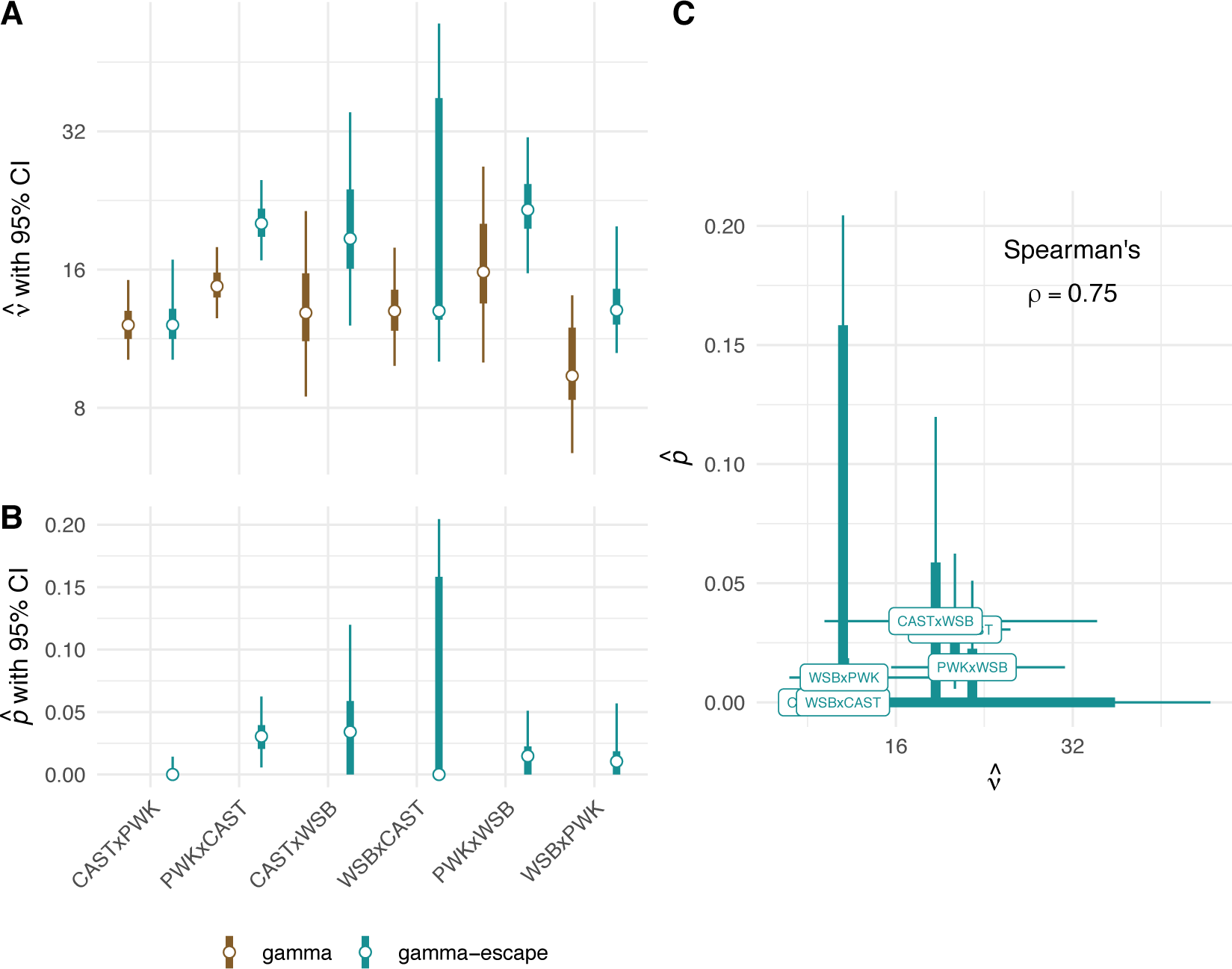
Interference parameters by genotype. (**A**) Estimates of strength of crossover interference by genotype, under the gamma and gamma-escape (Houseworth-Stahl) models. Error bars show 2.5% – 97.5% (thin) and 25% – 75% (thick) bootstrap confidence intervals. (**B**) Estimates of proportion of crossovers escaping interference by genotype. Error bars extend to the margin of the chart when confidence intervals include zero. (**C**) Apparent positive correlation between *p* and *ν* across genotypes.

**Table 6:** Comparison of gamma and gamma-escape models fit by maximum likelihood, allowing strength of crossover interference to vary by genotype (full) or not (null). df = degrees of freedom, LL = log-likelihood, AIC = Akaike information criterion

| model |  | df | LL | AIC |
| --- | --- | --- | --- | --- |
| gamma | null | 1 | -5514.8 | 11031.6 |
|  | full | 6 | -5507.0 | 11026.0 |
| gamma-escape | null | 2 | -5480.8 | 10965.5 |
|  | full | 12 | -5471.6 | 10967.3 |

**Table 7:** Pairwise comparisons between interference models. Entries above diagonal are test statistics, shown as “*χ*^2^ (df)”. Entries below diagnoal are *p*-values by likelihood ratio test. Blank cells indicate non-nested comparisons.

|  | gamma, null | gamma-escape, null | gamma, full | gamma-escape, full |
| --- | --- | --- | --- | --- |
| gamma, null | – | 68.1 (1) | 15.6 (5) | 86.3 (11) |
| gamma-escape, null | $1.6 \times 10^{-16}$ | – | – | 18.2 (10) |
| gamma, full | $8.2 \times 10^{-3}$ | – | – | 70.7 (6) |
| gamma-escape, full | $8.9 \times 10^{-14}$ | $5.1 \times 10^{-2}$ | $2.9 \times 10^{-13}$ | – |

We also discovered that it is challenging to estimate the proportion of crossovers that escape interference from this type of data. Intuitively we might expect that a wide range of (*ν, p*) pairs could be compatible with a given set of haplotype intervals – if a large number of crossovers escape interference (and can therefore be placed close together), then the strength of interference inferred from the remaining intercrossover distances will be higher. Inspection of (*ν, p*) pairs inferred from bootstrap replicates within genotypes confirms this suspicion (**Figure S9**). The positive correlation between rate of escape and strength of interference across samples from the same underlying distribution suggests non-identifiability in the statistical model.

### Strength of interference differs between some reciprocal crosses

Having provided evidence that interference strength varies across genotypes, we sought improved estimates of *ν* and associated uncertainties in order to make more detailed comparisons between genotypes. We developed a hierarchical Bayesian model that regularizes the *ν_i_* towards a common prior distribution, and also estimates a paternal age effect that is shared across genotypes. (We did not attempt to estimate interference escape in the Bayesian framework due to computational constraints and pathological behavior of MCMC with parameter correlations such as those noted in the previous section.) Because the method yields samples from the joint posterior distribution, it is straightforward to calculate derived quantities of interest such as groupwise comparisons. The Bayesian and maximum-likelihood methods provide very similar point estimates (root mean square difference = 0.51, or 5.3% of the smallest value of *ν_i_*), as shown in **Figure S6**.

Estimates of the strength of interference by genotype and paternal age are shown in **Figure 5**. Our estimates are close to the 11.3 obtained by Broman *et al*. (2002), the only other published application of the gamma model to house mice, although that study used an interspecies backcross between C57BL/6J and *Mus spretus*. The largest and smallest *ν_i_* values are obtained from the reciprocal crosses PWK/PhJ *×* WSB/EiJ and WSB/EiJ *×* PWK/PhJ, respectively. Contrasts between genotypes were calculated on the log scale (since *ν >* 0 by definition) and therefore represent ratios between values of *ν_i_* when transformed back to the natural scale (**Figure S7**). Three out of 15 comparisons reach conventional levels of statistical significance : [PWK/PhJ *×* WSB/EiJ vs WSB/EiJ *×* PWK/PhJ], [PWK/PhJ *×* CAST/EiJ vs WSB/EiJ *×* PWK/PhJ] and [CAST/EiJ *×* PWK/PhJ vs WSB/EiJ *×* PWK/PhJ]. Given the evidence for parent-of-origin effects on recombination rate, we focused particular attention on differences in strength of interference in reciprocal pairs (those with the same parents, but crossed in opposite directions). Interference is 1.62 times (95% HPDI 1.05 *—* 2.45 times) stronger in PWK/PhJ *×* WSB/EiJ than WSB/EiJ *×* PWK/PhJ; there is little difference between the other reciprocal pairs (**Figure S8**).

**Figure 5:**
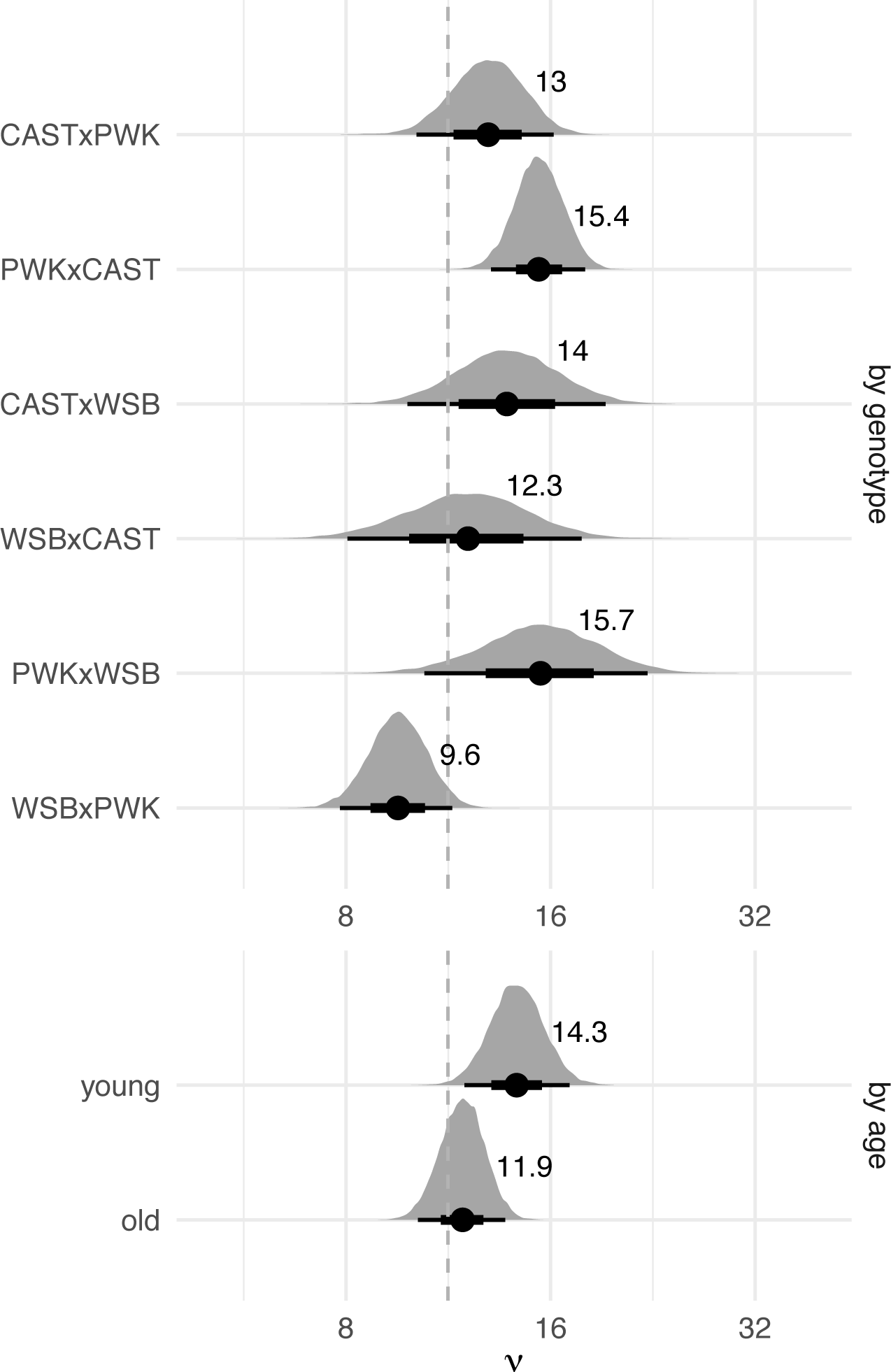
Posterior distributions of interference parameter (*ν*) by genotype, averaged over age groups; and by age group, averaged over genotypes. Solid dots and text labels show posterior mean; bars show 2.5% – 97.5% (thin) and 25% – 75% (thick) quantile intervals. Dashed line shows previous estimate from Broman *et al*. (2002). Note that the horizontal axis is in log scale.

Age affects the strength interference too, though less severely than genotype. Interference is 1.20 times (95% HPDI 0.97 *—* 1.49 times) stronger in younger versus older males. The null value (1.0) falls at the 5^th^ percentile of the posterior distribution.

It should be noted that the structure of the Bayesian model differs from that of the maximum-likelihood model in two important ways: first, by the inclusion of a term for the age effect; and second, by the fact that the *ν_i_* are estimated jointly in the Bayesian model, and separately within partitions of data in the maximum-likelihood model. We did also fit a null Bayesian model with a single value of *ν* and no age effect. The expected predictive accuracy (Vehtari *et al*. 2017) of the full and null models was very similar . However, we emphasize that the goal of the Bayesian analysis is parameter estimation rather than model selection.

## Discussion

Our portrait of male recombination obtained from crosses between wild-derived strains representing three subspecies of house mice corroborates and extends previous work in this area. We find that alleles from the *M. m. musculus* strain PWK/PhJ tend to increase recombination, whereas alleles from the *M. m. castaneus* strain CAST/EiJ tend to decrease it. This finding is consistent with autosomal QTL detected in CAST/EiJ *×* C57BL/6J (Murdoch *et al*. 2010) and CAST/EiJ *×* PWD/PhJ (Dumont and Payseur 2011b) intercrosses – QTL at which CAST/EiJ alleles reduce the number of crossovers. (PWD/PhJ is another *M. m. musculus* wild-derived strain that is very closely related to PWK/PhJ (Gregorová and Forejt 2000).) The pattern is also consistent with increases in recombination observed in consomic strains that carry certain autosomes from PWD/PhJ on the background of C57BL/6 (Balcova *et al*. 2016). In addition, autosomal alleles from PWK/PhJ are associated with more recombination, and those from CAST/EiJ are associated with less recombination, in the early generations of the Collaborative Cross (Liu *et al*. 2014), a panel of 8-way recombinant inbred lines. The decrease in recombination connected to PWK/PhJ mothers and the increase in recombination connected to CAST/EiJ mothers in this study also match previous inferences of X-linked QTL with effects in the same directions (Murdoch *et al*. 2010; Dumont and Payseur 2011b; Liu *et al*. 2014; Balcova *et al*. 2016).

Our results support the conclusion that crossover interference is strong in mice (Broman *et al*. 2002; de Boer *et al*. 2006). The shape parameter *ν* in the gamma model determines whether the distribution of distances between crossovers is exponential (*ν* = 1), corresponding to independence among crossovers, or humped (*ν >* 1), indicating that crossovers are spaced more regularly along the chromosome. Although values of *ν* are continuous, *ν —* 1 (when *nu* is an integer) is equivalent to the number of recombination intermediates that are located between two crossovers in the counting model of interference (Foss *et al*. 1993; Otto and Payseur 2019). Hence, our estimates of *ν* suggest that on average, crossovers are separated by 10 or more recombination intermediates that are repaired as non-crossovers in three subspecies of mice. Furthermore, the superior fit of the gamma-escape model to our data is in line with the existence of both interfering and non-interfering crossovers in mice (de Boer *et al*. 2006) as well as several other species (Otto and Payseur 2019).

The primary source of novelty in our study is the demonstration that the level of crossover interference varies among genotypes. Given that crosses were conducted in a common environment, heterogeneity among F1s detected using the gamma and gamma-escape models implies heritable differences in the level of interference and possibly the proportion of crossovers that avoid interference. The largest contrast comes from reciprocal crosses involving *M. m. musculus* (PWK/PhJ) and *M. m. domesticus* (WSB/EiJ). Although other maternal effects might contribute, stronger interference in PWK/PhJ *×* WSB/EiJ F1s than in WSB/EiJ *×* PWK/PhJ

F1s raises the possibility that the X chromosome modulates interference. This explanation is interesting in light of reports that PWK/PhJ and PWD/PhJ alleles on the X chromosome reduce recombination in other crosses (Dumont and Payseur 2011b; Liu *et al*. 2014), reductions that could be accomplished by amplifying interference. We also estimated the level of interference to be higher in PWK/PhJ *×* CAST/EiJ F1s than CAST/EiJ *×* PWK/PhJ F1s, though the difference is small. Another informative pairwise comparison is WSB/EiJ *×* PWK/PhJ F1s (lower interference) and CAST/EiJ *×* PWK/PhJ F1s (higher interference). These F1s differ by one set of autosomes as well as the X chromosome. This last comparison suggests that alleles from CAST/EiJ increase interference, again consistent in direction with the lower rate of recombination in CAST/EiJ than in WSB/EiJ and PWD/PhJ (Dumont and Payseur 2011a; Peterson and Payseur 2021). There is not a monotonic relationship between recombination rate and strength of interference among F1 genotypes (**Figure S10**). While correlations between the strength of interference and the rate of interference escape may be due in part to non-identifiability issues for the gamma-escape model, parameter estimates from the fit of this model might still contain biological information. For example, PWK/PhJ *×* CAST/EiJ F1s show higher levels of both interference and interference escape than CAST/EiJ *×* PWK/PhJ, while escape is similar in PWK/PhJ *×* WSB/EiJ and WSB/EiJ *×* PWK/PhJ despite significant difference in strength of interference. These differences warrant further investigation with methods that explicitly distinguish interfering from non-interfering crossovers based on their molecular features.

Although parameter estimates associated with each cross should be treated with some caution, our observation that the level of interference depends on genetic background provides evidence for evolution of this important determinant of crossover positioning. The three subspecies of mice to which WSB/EiJ, PWK/PhJ, and CAST/EiJ belong are estimated to have begun diverging about 350 000 *—* 500 000 years ago (reviewed in Phifer-Rixey and Nachman (2015)). To the extent that the wild-derived strains we examined are representative of their subspecies, this divergence time suggests the evolution of interference can occur over relatively short timescales.

The cellular and molecular causes of the variation we document are unclear. Multiple mechanisms of interference have been proposed (Otto and Payseur 2019). Recent discoveries in *Arabidopsis* species suggest that interference is created by diffusion-mediated growth of evenly spaced foci of the ubiquitin ligase HEI10 at the expense of closely spaced foci along the synaptonemal complex (Morgan *et al*. 2021; Durand *et al*. 2022; Fozard *et al*. 2023). Whether a similar “coarsening” mechanism operates in mice and contributes to the evolution of interference remains to be seen.

Another novel finding from our study is that older mouse males (at least 15 months of age) transmit an average of 0.61 more crossovers per meiosis than younger males (no more than 4 months of age). Although this effect is modest, it is consistent in magnitude and direction with a previous study in two classical inbred strains that found about 2 more interfering crossovers per spermatocyte in 1-year-old males versus males undergoing the first wave of spermatogenesis at 20 days postpartum (Vrooman *et al*. 2014). The effects of age on recombination rate in mammals are not well established in general. Studies reported no effect of paternal age in humans (Hussin *et al*. 2011) or sheep (Johnston *et al*. 2016). Maternal age has been associated with increased recombination in humans (Kong *et al*. 2004; Hussin *et al*. 2011; Campbell *et al*. 2015; Martin *et al*. 2015) and pigs (Lozada-Soto *et al*. 2021), decreased recombination in house mice (Polani and Jagiello 1976), increased and decreased recombination in cattle (Shen *et al*. 2021), and no change in recombination in sheep (Johnston *et al*. 2016). We also find that interference is about 9% stronger in older males, although the effect does not meet the nominal threshold for significance. Our results suggest that paternal age deserves further investigation as a modulator of crossover number and positioning.

We attach several caveats to our findings. First, distances between crossovers were computed by assuming that the standard male genetic map applies to all F1s, which could affect our characterization of interference. If the true recombination rate is higher in a given genotype than in the common map, we would expect more crossovers per interval than predicted by the map and downstream estimates of the strength of interference would be biased downward. Nevertheless, as shown in **Figure S10**, we do not find a consistent relationship between interference and recombination rate across genotypes. Second, our analyses are based on transmitted genotypes in live progeny, and therefore reflect the recombination landscape in viable gametes only. Certain crosses between *M. m. musculus* mothers and *M. m. domesticus* fathers generate F1 hybrid males with signs of reduced fertility caused by incompatibilities that disrupt meiosis I (Forejt and Iványi 1974; Forejt 1996; Good *et al*. 2008; Schwahn *et al*. 2018). If interference is systematically different in spermatocytes that are eliminated by meiotic checkpoints, or in gametes that do not mature into functional sperm, then our estimates of interference parameters will be biased. At this point there is no reason to suspect that the transmitted crossovers we surveyed comprise a non-random subset. Finally and most importantly, our study is restricted to males. Differences in meiosis between females and males in mice (Peterson and Payseur 2021) and other species (Basu-Roy *et al*. 2013; Campbell *et al*. 2016; Wang *et al*. 2016; Torgasheva and Borodin 2017) predict distinct evolutionary dynamics for interference in the two sexes.

## Acknowledgements

This project was launched in the laboratory of Fernando Pardo-Manuel de Villena in 2012 as an outgrowth of existing studies of aging and male infertility. Support was provided by the following grants from the National Institutes of Health: F30 MH103925 (Andrew Morgan), K01 MH094406 (James J Crowley), R21 MH096261 (Fernando Pardo-Manuel de Villena), R01 HD065024 (Fernando Pardo-Manuel de Villena), P50 MH090338 (Fernando Pardo-Manuel de Villena), P50 HG006582 (Fernando Pardo-Manuel de Villena), U42 OD010924 (Terry Magnuson), and R35 GM139412 (Bret Payseur). We thank Timothy A Bell and James J Crowley for mouse husbandry. We thank Grace Clark, Justin Gooch, the late Mark Calaway, and Darla Miller for assistance with DNA preparation and submission of samples for genotyping. We benefited from fruitful correspondence with Fernando Pardo-Manuel de Villena, Beth Dumont, Karl Broman and several other members of the mouse genetics community in the course of this work.

## Data availability

Sample metadata, raw and processed genotype data are available on Figshare: doi:10.6084/m9.figshare.25859950. Analysis code is available on Github: https://github.com/andrewparkermorgan/mouse_crossover_interference.

## Author contributions

APM designed and performed the analyses. APM and BAP and wrote the manuscript.

## Supplementary material

**Table S1.** Sample metadata for *n* = 503 N2 progeny of F1 fathers.

**Table S2.** All inferred autosomal haplotype segments (*m* = 15 744) for *n* = 503 mice.

**Table S3.** Number of autosomal crossovers inferred per individual for *n* = 503 mice.

Raw and processed genotype matrices are available from Figshare.

**Figure S1:**
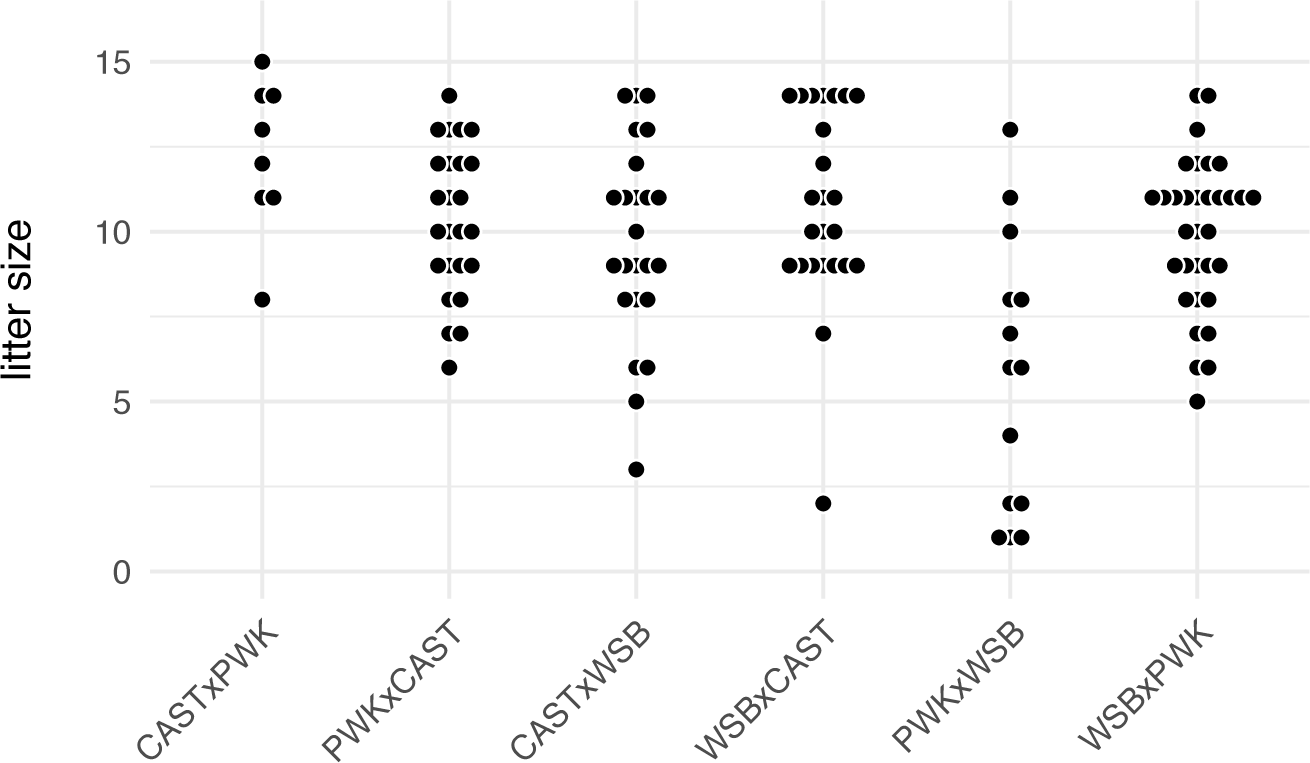
Litter size by sire genotype

**Figure S2:**
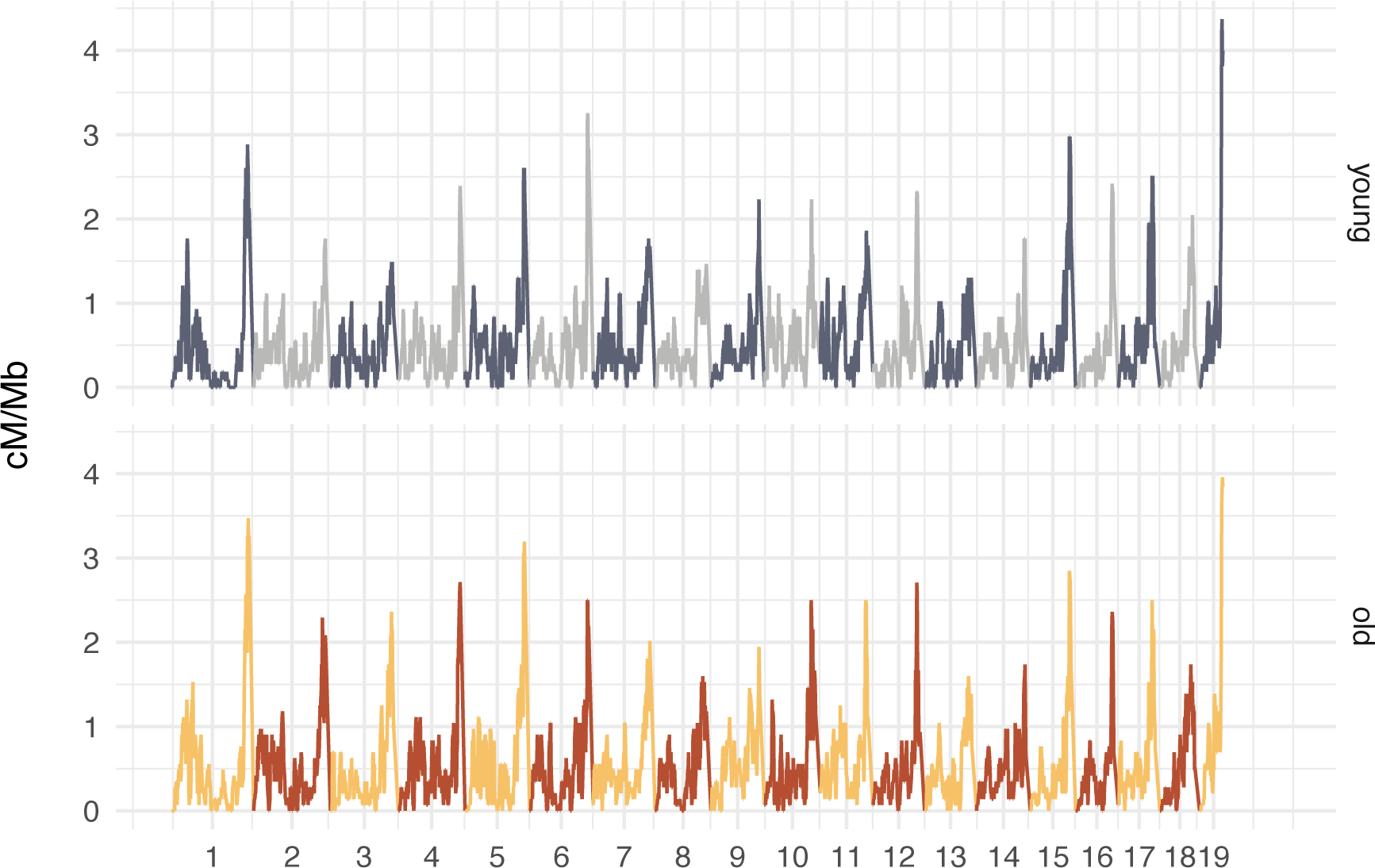
Recombination rate (in cM/Mb) in 5 Mb sliding windows across the autosomes, estimated separately for young and old males.

**Figure S3:**
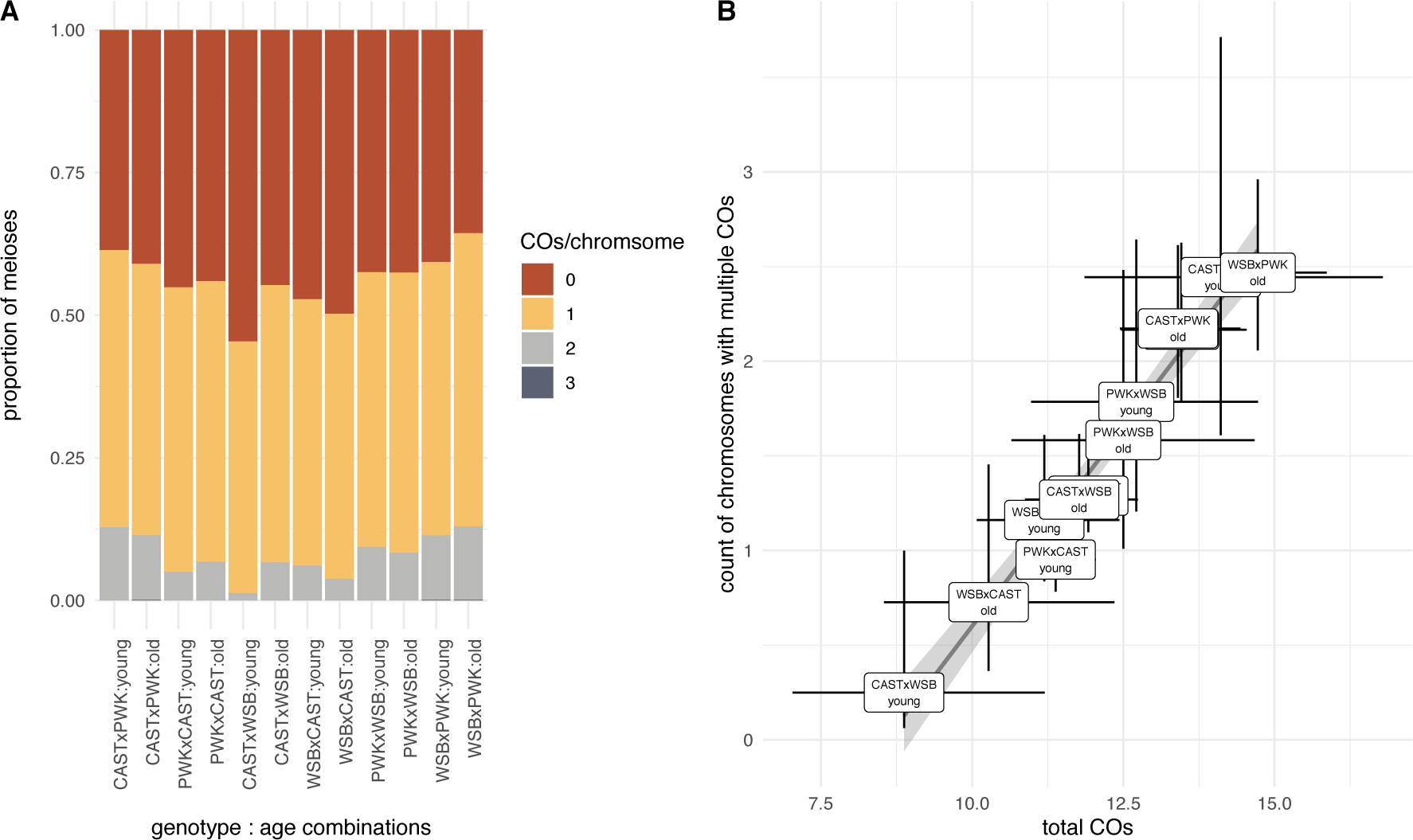
The number of multiply-recombinant chromosomes increases smoothly with recombination rate. (**A**) Distribution of chromosomes by number of crossovers in each genotype-by-age combination. (**B**) Count of chromosomes with multiple crossovers versus total number of crossovers in each genotype-by-age combination. Error bars are 95% confidence intervals from generalized linear model with genotype, age and their interaction as covariates. Solid grey line is a linear fit through the group means.

**Figure S4:**
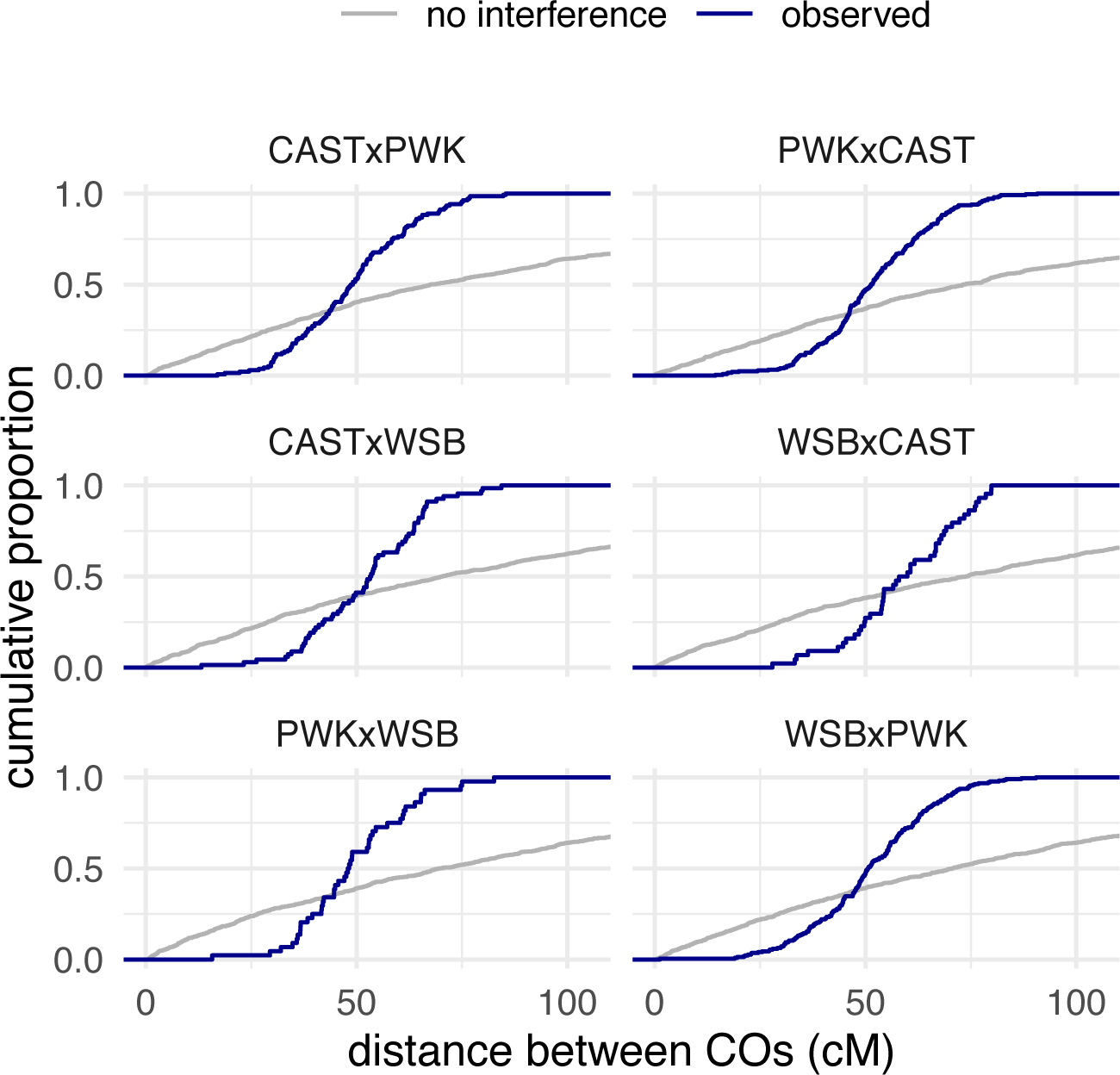
Empirical cumulative distribution of distance between crossovers (COs) on the same chromosome (blue curves), compared to expected distribution without crossover interference.

**Figure S5:**
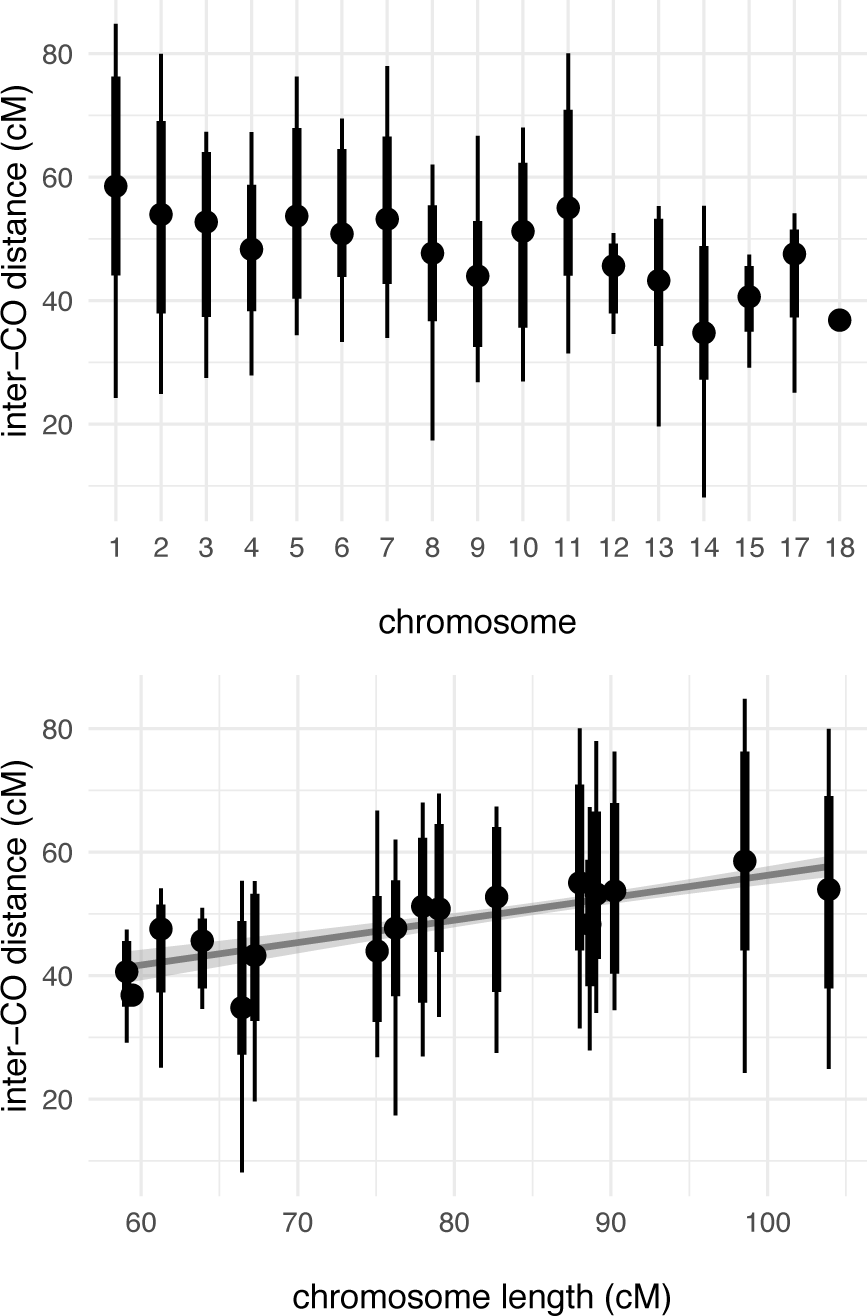
Distribution of inter-crossover distance by chromosome, for chromosomes on which multiply-recombinant progeny were obtained, plotted by chromosome number (**A**) and against total chromosome length (**B**). Solid dots are medians; bars show 2.5% – 97.5% (thin) and 25% – 75% (thick) quantile intervals. Solid grey line is a linear fit through the per-chromosome medians.

**Figure S6:**
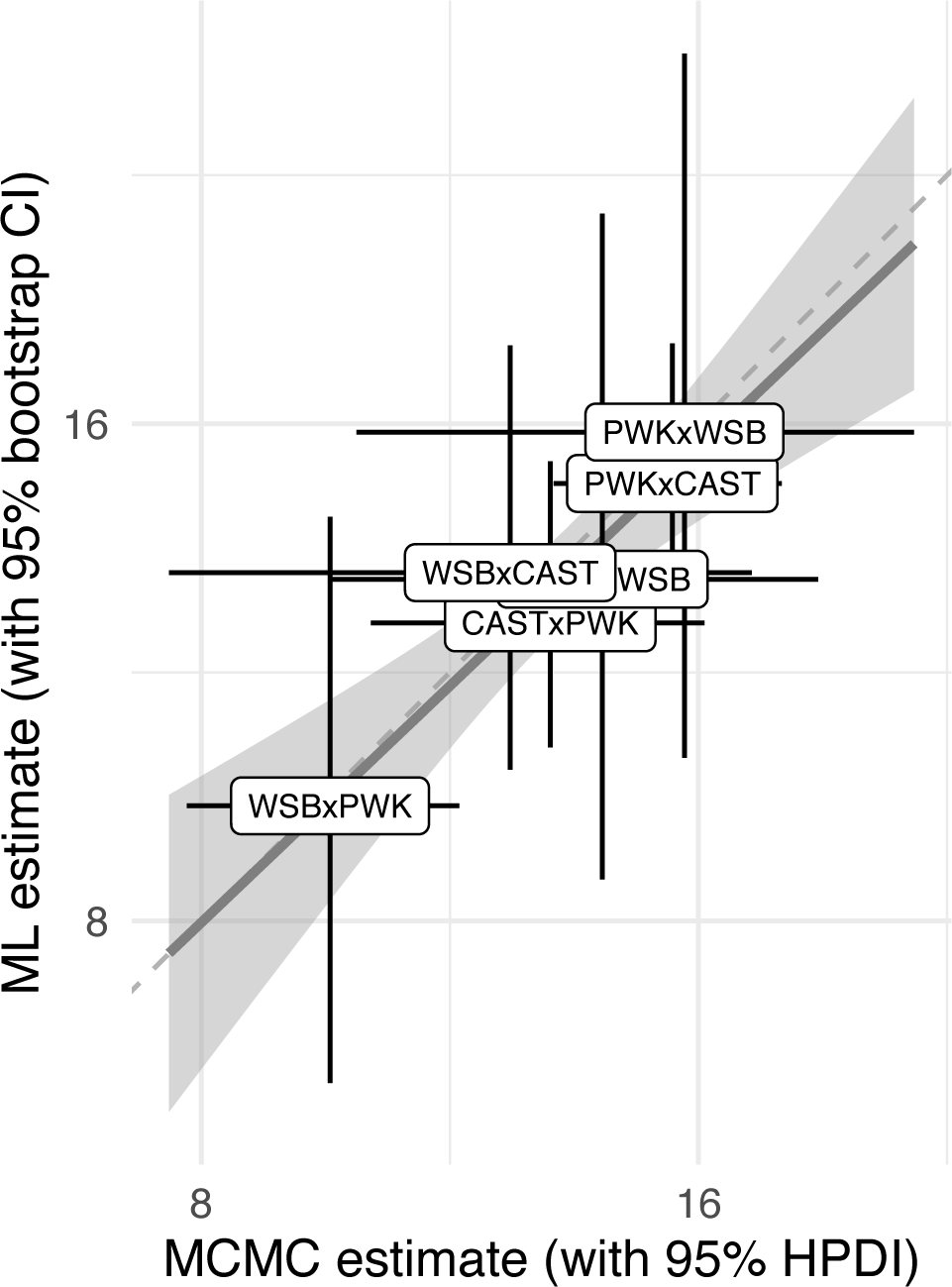
Comparison of group-wise estimates of the interference parameter *ν* obtained by maximum-likelihood versus joint estimates from Bayesian hierarchical model by MCMC. Solid grey line is a linear fit through the point estimates; dashed grey line has unit slope and passes through the origin.

**Figure S7:**
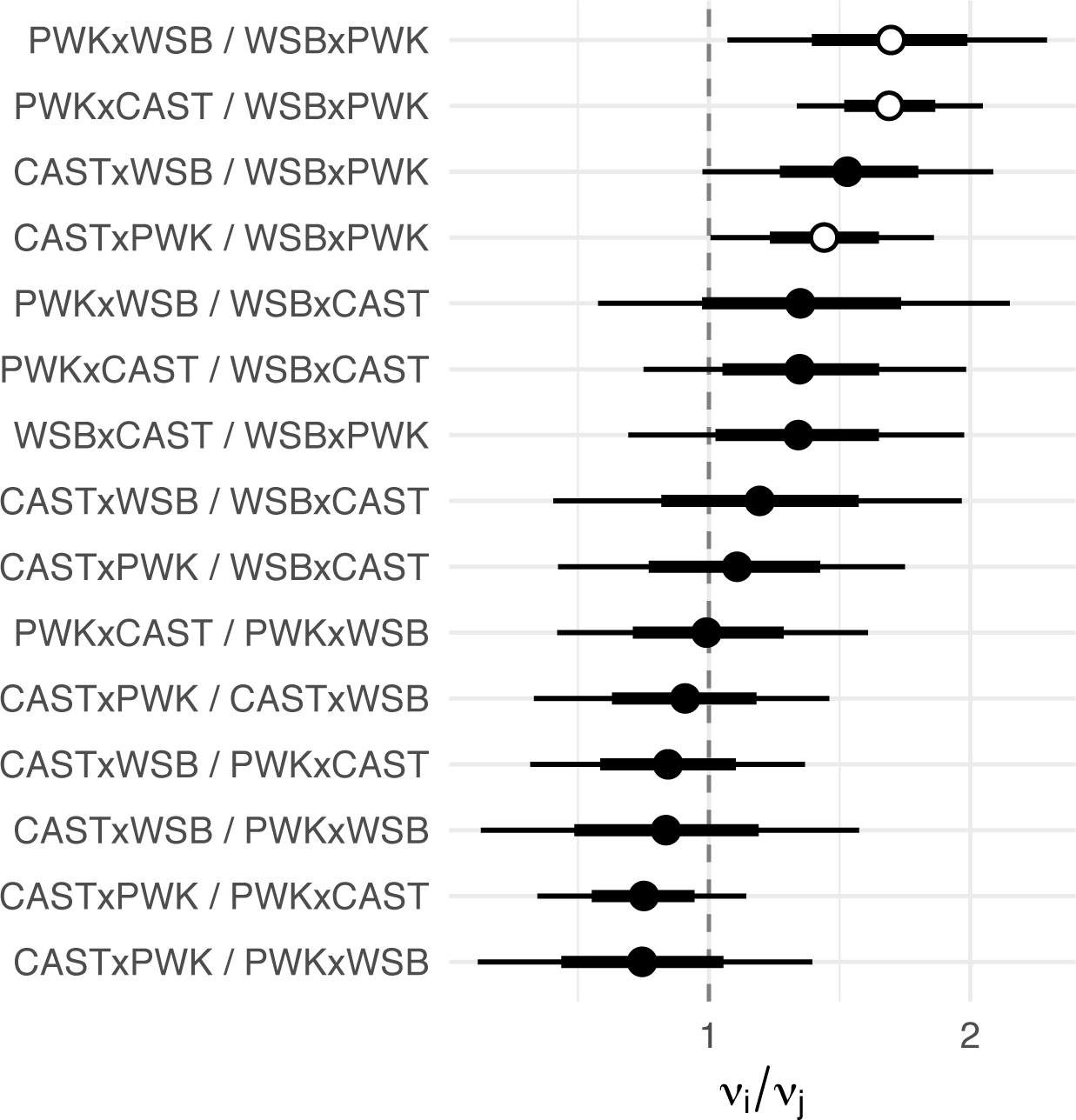
All pairwise contrasts between per-genotype estimates of strength of interference. Dots show posterior means and bars show 2.5% – 97.5% (thin) and 25% – 75% (thick) quantile intervals. Open dots indicate comparisons that meet nominal statistical significance; filled dots nominally non-significant. Dashed line gives the null value of 1. Note that the horizontal axis is in log scale.

**Figure S8:**
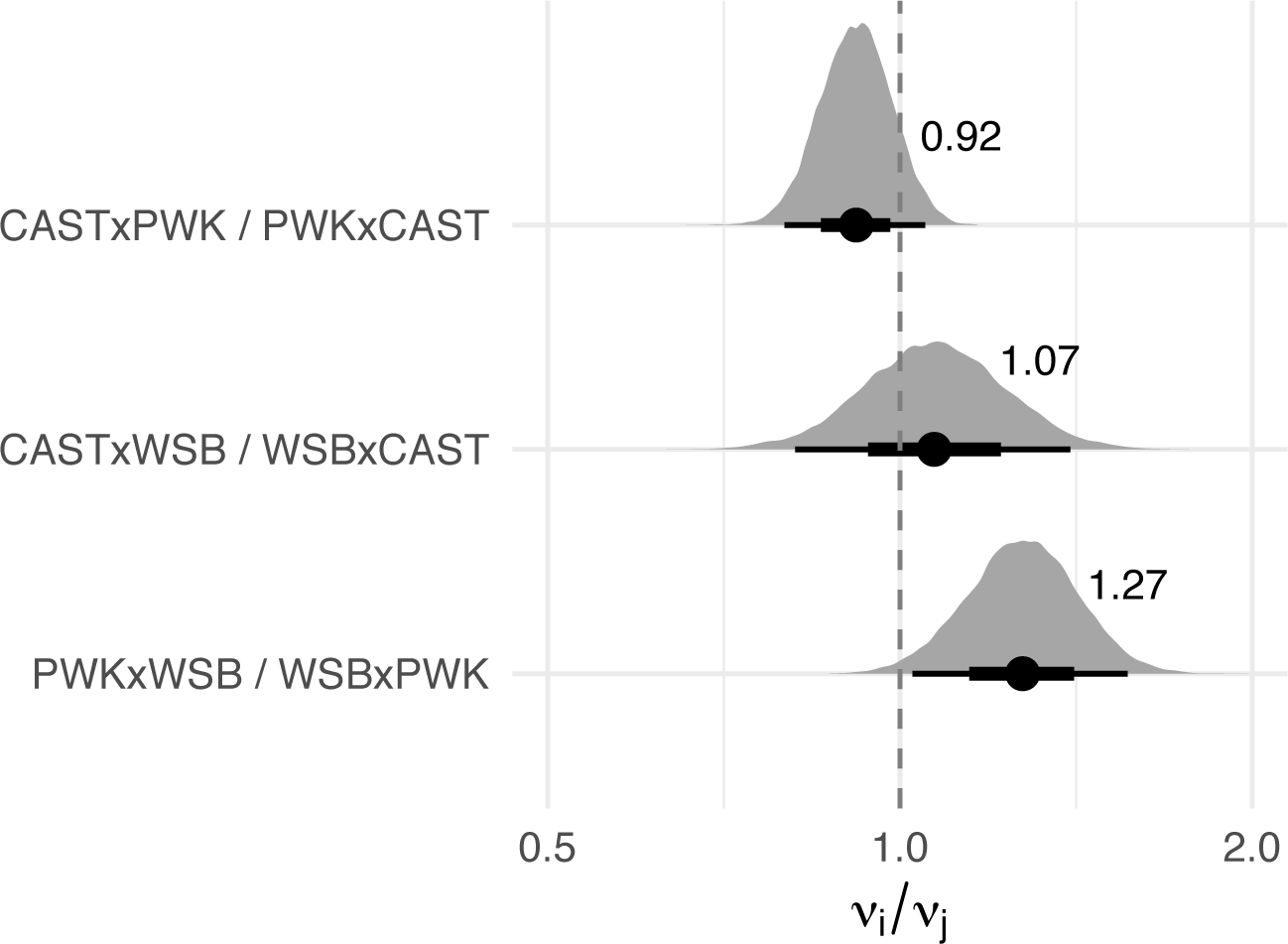
Contrasts between estimates of strength of interference in reciprocal genotypes. Text labels show posterior means. Dashed line gives the null value of 1. Note that the horizontal axis is in log scale.

**Figure S9:**
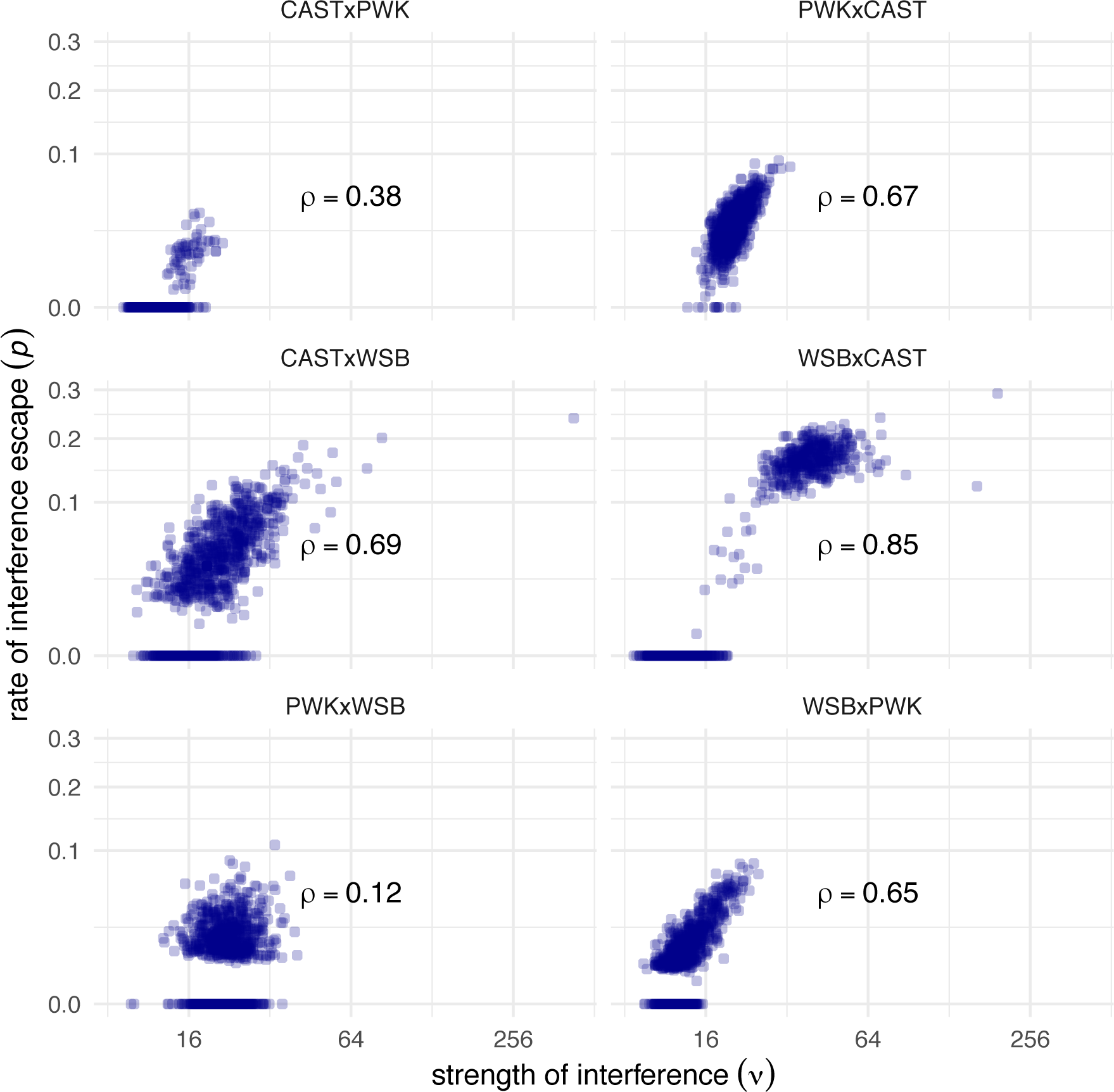
Maximum-likelihood estimates for the strength of interference (*ν*) and proportion of non-interfering crossovers (*p*) for 1000 bootstrap replicates of crossover data from six genotypes. Note that horizontal axis is in log scale and vertical axis in square-root scale.

**Figure S10:**
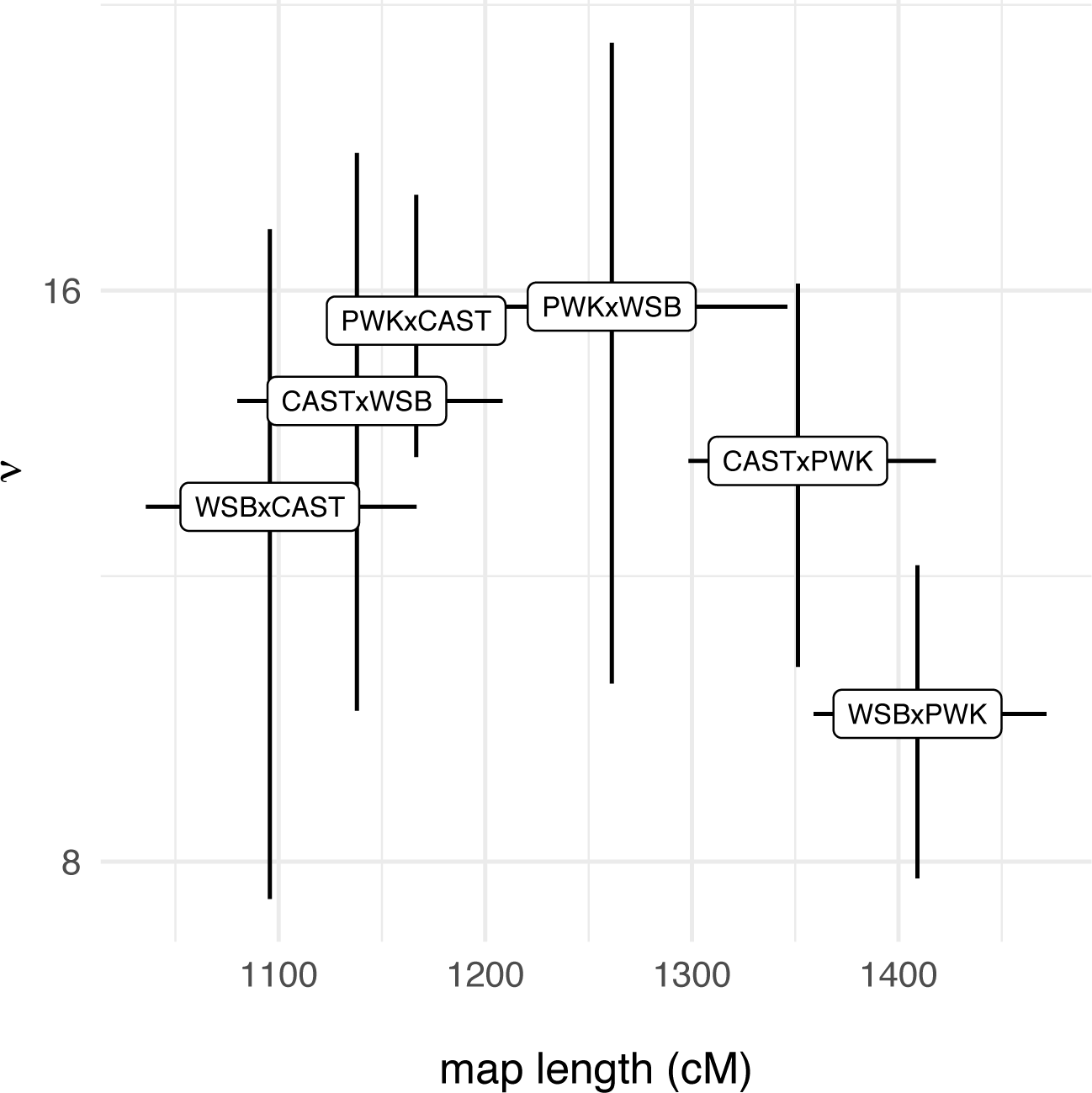
Relationship between length of the recombination map (in cM) and strength of crossover interference (*ν*). Error bars give 95% confidence bounds, calculated as described for the respective parameters elsewhere.

